# Development and validation of an experimental life support system to study the impact of ultraviolet B radiation and temperature on coral reef microbial communities

**DOI:** 10.1101/2023.03.01.530425

**Authors:** T.M. Stuij, D.F.R. Cleary, R.J.M. Rocha, A.R.M. Polonia, D.A.M. Silva, J.C. Frommlet, A. Louvado, Y. M. Huang, N. Van der Windt, N.J. De Voogd, N.C.M. Gomes

## Abstract

In the present study, we developed and validated an experimental life support system (ELSS) designed to investigate the response of coral reef associated bacterial communities to increases in temperature and UVB intensity. The ELSS consisted of 32 independent microcosms, which enables researchers to study the individual and interactive effects of up to three factors using a full factorial experimental design. Temperature can be controlled using water-baths. UV exposure was introduced to the system using UV fluorescent lights. Individual UVB-opaque polyester films were added to the microcosms using a random design. In the validation experiment (stable temperature and no UVB), a coral reef environment was simulated using a layer of coral reef sediment, synthetic seawater, and specimens from five benthic reef species. The species used were two hard corals *Montipora digitata* and *Montipora capricornis*, a soft coral *Sarcophyton glaucum*, a zoanthid *Zoanthus* sp., and a sponge *Chondrilla* sp.. To validate the system, we assessed physical and chemical parameters and characterised host and free-living bacterial communities of the ELSS over 34 days and compared these data to those observed in natural reef ecosystems. Water temperature, dissolved oxygen, pH, salinity and dissolved nutrients in the ELSS were similar to those at shallow coral reef sites. Sediment bacterial diversity and composition were more similar to natural-type communities at day 29 and 34 than at day 8 after transfer to the microcosms, indicating a return to natural-type conditions following an initial, apparent perturbation phase. Transplantation significantly altered the bacterial community composition of *M. digitata* and *Chondrilla* sp. and increased coral photosynthetic efficiency compared to before transplantation. These results highlight the importance of *M. digitata* and *Chondrilla* sp. microbiomes to host adaptation following potential stress events.. Altogether, our results validated the suitability of the ELLS developed in this study as a model system to investigate the responses of coral reef associated bacterial communities to shifts of temperature and UVB radiation and potentially other environmental conditions (e.g., environmental pollution).

## 1. Introduction

Coral reefs are among the most diverse and productive of marine ecosystems and are home to an estimated 25% of known marine species (Fisher et al., 2015). They also provide important economic services including coastal protection, tourism and fisheries (Fisher et al., 2015; Moberg and Folke, 1999; Plaisance et al., 2011). Natural environmental perturbations, such as storms and changes in sea levels, have always strongly affected coral reefs, but anthropogenic perturbations have clearly and increasingly affected the health and resilience of coral reefs (Baker et al., 2008; De’ath et al., 2012; Glynn et al., 2017). For example, the frequency of extreme El Niño southern oscillation (ENSO) events has increased over the past decades (Cai et al., 2015; Wang et al., 2019; Ying et al., 2022); this has been linked to an increase in larger-scale and more severe coral bleaching events (Baker et al., 2008; Eakin et al., 2019; McGowan and Theobald, 2017). Given the above, it is important to understand how coral reef organisms respond to environmental perturbations, particularly those related to predicted increases in temperature and UVB intensity and their relationship with large-scale processes (Masson-Delmotte et al., 2021).

Observational studies in coral reef ecosystems have provided important insights into the relationship between coral bleaching and temperature (Hughes et al., 2017), water turbidity (Sully and van Woesik, 2020), and large-scale ocean-atmospheric phenomena such as ENSO events (Eakin et al., 2019; McGowan and Theobald, 2017; Zhang et al., 2017). However, coral reefs are highly complex ecosystems, and the direct and indirect impact of multiple environmental parameters on community dynamics complicate the ability to delineate correlation from causation (Dizon and Yap, 2006). In contrast to *in situ* studies of natural habitats, randomised controlled microcosm experiments provide a means of reducing the environmental complexity and create controlled conditions, under which the effects of specific factors can be assessed. This enables hypothesis testing, and in the case of multifactorial experiments, testing for independent and interactive effects of different environmental factors (Coelho et al., 2015, 2013; Findlay et al., 2008; Schubert and Wilke, 2017). This facilitates a more mechanistic understanding of the relationship between specific environmental factors and the response variable or variables of interest (Boyd et al., 2018; Coelho et al., 2015, 2013). Moreover, microcosm-based studies offer a way to study organismal responses to specific and controllable treatments, which may not be possible in a natural setting. For example, previous microcosm studies revealed the interactive effect of nutrient enrichment, temperature and UV radiation on coral physiology (Courtial et al., 2017; Schlöder and D’Croz, 2004).

In the last years a growing body of studies have highlighted the importance of reef microbial communities for coral reef resistance and resilience to environmental perturbations (Ainsworth et al., 2010; Glasl et al., 2019; van Oppen and Blackall, 2019; Vanwonterghem and Webster, 2020). These communities play important roles in biogeochemical cycling, degradation of pollutants, coral nutrition and defense against pathogens and predators (Alongi, 1994; Dong et al., 2022; Rädecker et al., 2015; Ricci et al., 2019; Rosenberg et al., 2007; Wild et al., 2004). Both non-host (e.g., in sediment and seawater) and host-associated communities, have been shown to respond to spatial and environmental processes including large and local-scale perturbations (Coelho et al., 2015; Fan et al., 2013; McDevitt-Irwin et al., 2017). In corals, elevated temperature and UVB in combination with water pollution and overfishing were shown to adversely affect putative symbiotic bacteria (e.g., *Endozoicomonas*) and increase the abundance of opportunistic bacteria (e.g., *Vibrio* species) (McDevitt-Irwin et al., 2017). Moreover, compositional shifts in sponge bacterial communities following environmental perturbations coincided with a reduced abundance of symbiotic functions (Fan et al., 2013; Pita et al., 2018; Vanwonterghem and Webster, 2020).

In the current study, we developed and validated an experimental life support system (ELSS), designed with the purpose to investigate the response of coral reef associated bacterial communities to increases in temperature and UVB intensity. We monitored physical and chemical parameters, assessed coral photosynthetic activity and investigated sediment, water, and host-associated bacterial communities. Subsequently, we validated the system by comparing the conditions and bacterial communities in the ELSS with those occurring in the natural coral reef environments.

## 2. Methods

### 2.1. ELSS design

The design of the ELSS developed in this study was based on a microcosm system previously developed to assess the effects of global climate change and environmental contamination on sediment communities (Coelho et al., 2013). The ELSS includes two frames of 16 microcosms (32 in total) and was shown to replicate fundamental aspects of biological activity in sediments of coastal marine and estuarine environments. For the current study, this system was modified to allow the evaluation of the effects of UV and temperature on coral reefs under laboratory-controlled conditions. The 32 microcosms (glass aquaria, 23 cm in height, 16 cm length and 12 cm width) were each connected to another aquarium (referred to as reservoir, 30 cm in height, 12 cm length and 12 cm width), see Fig. 1. The microcosms and reservoirs contained outflow-holes (two centimetres in diameter) positioned 3 and 5 cm below the top of the glass, respectively. Each reservoir was equipped with a small hydraulic pump (compactON 300, EHEIM) to pump water out of the microcosm into the reservoir, where subsequently the water level would rise and flow out into the accompanied microcosm with a constant flow rate of approximately 8.64 ml/s (s = 0.66 ml/s). Each reservoir-microcosm unit contained a functional water volume of approximately 5 L.

**Fig. 1.**
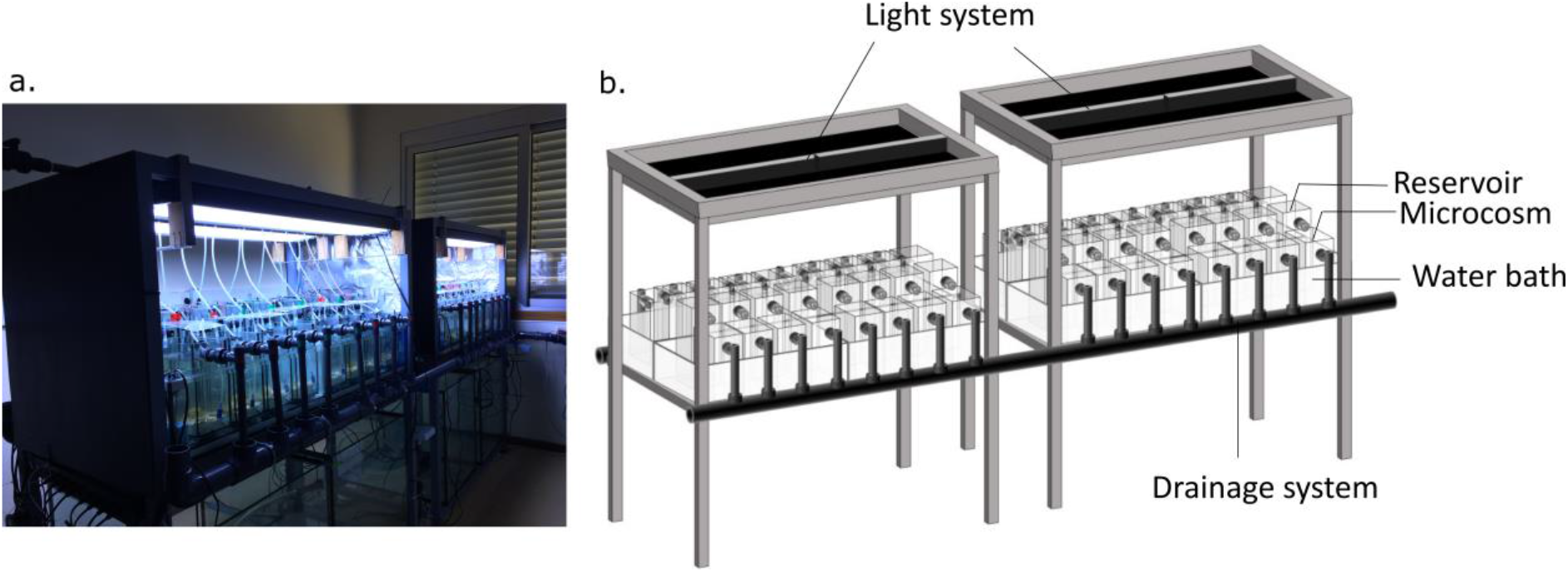
Picture (a) and graphical representation (b) of the experimental life support system.

### 2.2. Water changes, temperature, light and aeration control system

A semi-static setup was chosen as it offered the possibility to have constant flow (water aeration and circulation) into the microcosms along with water renewal and temperature homogenization. Water was partly renewed daily by replacing 1 L of water with newly prepared synthetic seawater. Synthetic seawater was prepared by mixing coral reef salt (including associated oligonutrients) (CORAL PRO SALT, Red sea) with deionized water (produced in a four-stage reverse osmosis unit - V2Pure 360) to a concentration of 35 ppt. To obtain sufficient mixing, the synthetic seawater was prepared in a large container supplied with a recirculating pump at least 12 hours prior to addition to the reservoirs.

To minimize temperature fluctuations in the microcosms, temperature was regulated using water bath tanks, each of which surrounded four microcosms (Fig. 1, Coelho et al. 2013). The temperature in these tanks was regulated by water heaters (V2Therm 100 Digital heater) equipped with an internal thermostat. For 34 days, water temperature was kept at 28 ⁰C. The modular design of the ELSS set up enables randomized split-plot design experiments, whereby each plot consists of four microcosms of equal temperature.

Lighting was controlled by four fully programmable luminaire systems (Reef - SET, Rees, Germany), each holding eight fluorescent lamps (Coelho et al., 2013). During the current experiment, four UV fluorescent tubes (SolarRaptor, T5/54W) and four full spectra fluorescent tubes (ATI AquaBlue Special, T5/54W) were connected alternately. To simulate photoperiod conditions of tropical latitudes, the lamps were programmed to a 12 h diurnal light cycle with light intensity varying from 2.61*10^−3^ J cm^−2^ s^−1^ in the morning to 8.82*10^−3^ J cm^−2^ s^−1^ at mid-day (measured using a Fibre optic probe positioned at the water surface, Flame spectrometer, Ocean Optics). The total light energy transmitted during the day equalled 256 J cm^−2^ day^−1^, of which 97.7% came from the photoactive radiation (PAR) wavelengths (300 – 700), 1.85% from the UVA wavelengths (315 – 400) and 0.43% from the UVB wavelengths (280 – 315). Photosynthetic photon flux density (PPFD, mol photons m^−2^ s^−1^) at the sediment-water interface was measured using an underwater PAR meter (MQ-510 PAR/Quantum-Meter Underwater, Apogee, USA). A total of 3.47 ± 0.68 mol photons m^−2^ d^−1^ reached the sediment surface. A transparent polyester film (Folanorm SF-AS, Folex coating, Köln, Germany) was used to block UVB light (290 – 320 nm) (Müller et al., 2009; Zacher et al., 2007). This film absorbed 90% of the UVB irradiance, 31% of UVA and 9% of PAR irradiance. A detailed figure of the light spectrum can be found in Supplementary Fig. 1. When the ELSS is used to investigate the effects of UVB on coral reef organisms, this film can be removed from the microcosms. All microcosms were connected to an air pump (530L/h, Resun) providing equal and constant aeration in each microcosm via a small hose and a diffuser stone (1.5 cm in diameter, 3 cm in length).

### 2.3. Coral reef sediment and organisms

The bottom of each microcosm received a sediment layer of ∼3 cm, equivalent to 700 g wet weight per microcosm. This sediment consisted of a mixture of commercially available (Reef Pink dry aragonite sand, Red Sea) and natural coral reef sediment. Natural sediment was collected from a coral reef south of Fongguei, Penghu Islands, Taiwan (22°19’50.5”N 120°22’19.8”E). The sediment was transported to our laboratory in Portugal in a coolbox and stored at 4 ⁰C until use. This transportation between collection and the microcosm experiment occurred in the minimal amount of time possible (5 days). The commercial sediment was washed and sterilized. Sterilization was performed 3 times by autoclavation at 121°C for 20 minutes (Otte et al., 2018). 20 kg of sterilized commercial sediment was spiked with 4 kgs of natural coral reef sediment, which equates to a 1 to 6 ratio of live to sterilized sediment. The mixture was thoroughly homogenized and added to the microcosms.

The ELSS was left to stabilize with spiked reef sediment and synthetic seawater for the first eight days of the experiment. Subsequently, five reef animal species were added, after which the system ran continuously for an additional 26 days. Overall, the system ran continuously for 34 days. The species used in the ELSS included two hard corals, *Montipora digitata* (Dana, 1846) and *Montipora capricornis* Veron 1985, one soft coral, *Sarcophyton glaucum* (Quoy & Gaimard, 1833), one zoanthid, *Zoanthus* sp., and one sponge, *Chondrilla* sp.. All animals used in this study were previously grown in aquaria at ECOMARE (CESAM, University of Aveiro, Portugal) (detailed explanation the fragmentation process of the organisms can be found in the Supplementary material, Rocha et al. 2015).

### 2.4. Physical and chemical analysis

Water quality was monitored daily by measuring temperature, pH, dissolved oxygen and salinity (Multi 3420 multimeter, WTW GmbH, Weilheim, Germany). Samples for determining dissolved inorganic nutrient concentrations (nitrate NO^3-^; nitrite NO^2-^ ammonium NH^4+^, and phosphate PO_4_^3-^) in the water column were taken every 7 days, using disposable syringes (50 ml), and were measured immediately, according to the Sali test kit protocol (colorimetric method test kit, salifert, Aquarium Masters).

In addition to the water column nutrients, we analysed dissolved inorganic nutrients (nitrate NO^3-^; nitrite NO^2-^; ammonium NH^4+^; phosphate PO_4_^3-^; and sulphate SO_4_^2-^) and total organic carbon (TOC) concentrations from samples of sediment porewater after eight, 28 and 34 days. Samples were obtained using Rhizon flex samplers with a pore size of 0.6 μm (product number 19.60.25F, Rhizosphere Research Products, Wageningen, The Netherlands). To collect the sample, the sampler was connected to 50 ml disposable syringes. Nitrate NO^3-^; nitrite NO^2-^; ammonium NH^4+^; and phosphate PO_4_^3-^were analysed on the day of sampling using photometric methods following the standard analytical protocol of the Supelco Spectroquant® test kits 1.14942, 1.14776, 1.14752 and 1.14848, respectively (Merck, Darmstadt, Germany). From the obtained absorption value, nutrient concentrations were calculated using standard curves, which were determined from standard solutions prior to sample analysis. Immediately after collection, aliquots of the pore water samples were sent to an external service provider (A3lab, Ilhavo, Portugal), for sulphate SO_4_^2-^and TOC analysis. SO_4_^2-^was measured by turbidimetry using discrete spectrophotometry following the standard analytical method: CZ_SOP_D06_02_016 (SM 4500-SO4). TOC was measured using infra-red (IR) detection following the standard analytical method: CZ_SOP_D06_02_056 (SM 5310).

### 2.5. In vivo chlorophyll fluorescence analysis

Chlorophyll fluorescence of the corals and zoanthid was measured in vivo using a pulse amplitude modulation (PAM) fluorometer (Walz™), before adding the specimens to the microcosms and at the end of the experiment. Fluorescence was measured in dark-adapted samples (for 20 min), with Junior PAM and WinControl3 software (Walz™). Saturating light pulses (450 nm) were performed perpendicularly to the sample surface, with a 1.5 mm fibre optic. The maximum quantum yield (Fv/Fm) of photosystem II was calculated as 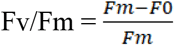.

### 2.6. Bacterial community analysis

#### 2.6.1. Sampling and DNA extraction

Sediment, water and porewater samples were collected after 8, 29 and 34 days of four independent microcosms. Reef animals were sampled at the end of the experiment (after 34 days) and rinsed with filtered (0.22-μm pore size) synthetic seawater to remove loosely attached organisms. Additionally, samples from the natural sediment collected in Taiwan and from benthic macro-organisms prior to addition to the microcosms (hereafter referred to as “Pre”) were preserved for further analysis (see fig. 2). During sediment collection in Taiwan, three sediment samples of approximately 20 g were collected *in situ* and immediately preserved in 96% alcohol. From the microcosms, a composite sediment sample consisting of four smaller subsamples (∼3 g each) was obtained using a sterilized scoop. The subsamples were taken haphazardly from each microcosm (1 cm of surface sediment with a ∼ 2 cm diameter). One sample from the sterilized commercial sediment was obtained as control for ELSS contamination with environmental DNA (Torti et al., 2015) and sample collection (Hornung et al., 2019).

**Fig. 2.**
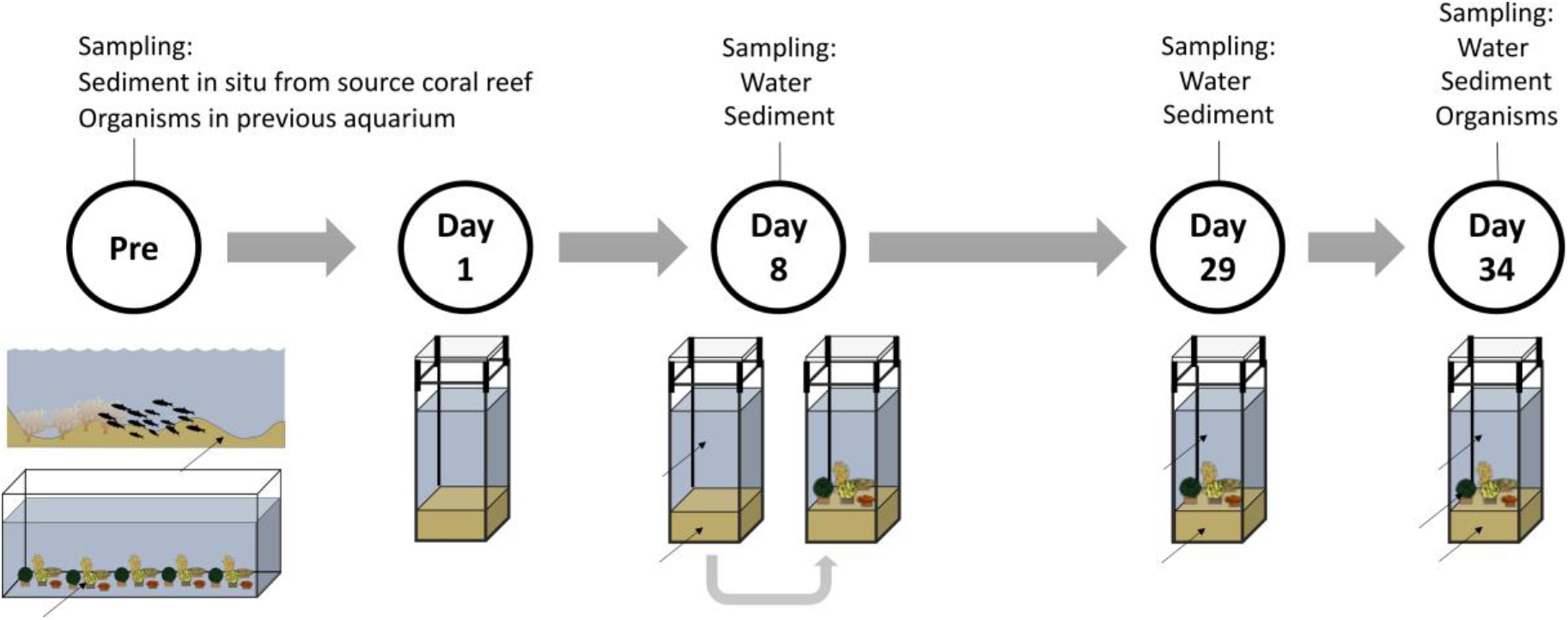
Graphical summary of the samples collected to study bacterial communities.

Water was sampled by filtering 250 ml of water over a Millipore® Isopore polycarbonate membrane filter (0.22-μm pore size; Millipore®) using a vacuum filtration system. All sediment samples, whole membrane filters and reef organisms were frozen at -80 ⁰C until DNA extraction.

PCR-ready genomic DNA was isolated from all samples using the FastDNA® SPIN soil Kit (MPbiomedicals) following the manufacturer’s instructions. Briefly, the whole membrane filter (bacterioplankton communities) and ± 500 mg of sediment and host organisms were transferred to Lysing Matrix E tubes containing a mixture of ceramic and silica particles. Due to the small size of *Zoanthus* sp. (< 60 mg) and *Chondrilla* sp. (varied between 130 and 500 mg), whole organisms were used for extraction. For *M. digitata* and *M. capricornis*, fragments (tissue and skeleton) were first snap frozen in liquid nitrogen and subsequently grinded using mortar and pestle. Blank negative controls, in which either no tissue or a sterile membrane filter was added to the Lysing Matrix E tubes, were also included. Microbial cell lysis was performed in the FastPrep Instrument (MP biomedicals) for 2 × 40 seconds at a speed setting of 6.0 ms-1. Extracted DNA was eluted in 50 μl of DNase/pyrogen-free water and stored at -20 ⁰C until further use.

#### 2.6.2. 16S rRNA gene library preparation and sequencing

The V3/V4 variable region of the 16S rRNA gene was amplified using primers 341F 5’CCTACGGGNGGCWGCAG′3 and 785R 5′GACTACHVGGGTATCTAATCC′3 (Klindworth et al., 2013 with Illumina Nextera XT overhang adapters for a dual-barcoding PCR library preparation approach. First PCRs were performed using 1 to 3 μl of DNA template, 10 μl of HS ReadyMix (KAPA HiFi Roche), and 0.6 μl of the forward and reverse primers in a concentration of 10 pmol^−1^μl^−1^. Reaction mixes were finalized by the addition of mQ water (Ultrapure) to a final volume of 20 μl. The PCR conditions consisted of initial denaturing at 95 °C for 3 min, followed by 30 cycles of 98 °C for 20 s, 57 °C for 30 s, and 72 °C for 30 s, after which a final elongation step at 72 °C for 10 min was performed. We checked for the success of amplification, relative intensity of the bands, and contamination using 2% Invitrogen E-gels with 3 μl of PCR product.

Subsequently, PCR products were cleaned with magnetic beads at a ratio of 0.9:1 using a magnetic extractor stamp, after which a second PCR was performed. The 25 μl reaction mix consisted of 4 μl of the first PCR product, 12.5 μl HS ReadyMix (KAPA HiFi Roche), 2 × 1 μl (concentration of 10pmol/μl) MiSeq Nextera XT adapters (dual indexed, Illumina), and 6.5 μl of mQ water (Ultrapure). PCR conditions consisted of initial denaturing at 95 °C for 3 min, followed by 8 cycles of 98 °C for 20 s, 55 °C for 30 s, and 72 °C for 30 s, after which a final elongation step at 72 °C for 5 min was performed. DNA molarity and fragment sizes of the resulted PCR products were measured on a fragment analyzer 5300 (Agilent) and subsequently, each 96-well plate was normalized and pooled together in subpools using the Qiagen QIAgility. The DNA molarity of the subpools was measured on the Agilent 4150 TapeStation to combine the subpools into a final pool. The resulted pool of normalized DNA was cleaned one last time using magnetic beads at a ratio of 0.65:1 and thereafter sequenced at a commercial company (Baseclear, Leiden, The Netherlands) on the Illumina MiSeq platform using 2 × 300 bp paired-end sequencing (Illumina MiSeq PE300). Three negative control samples were included to detect possible contamination during library preparation and sequencing. Sequences from each end were paired following Q25 quality trimming and removal of short reads (<150 bp). The DNA sequences generated in this study can be downloaded from NCBI BioProject Id PRJNA904682.

The 16S rRNA gene amplicon libraries were imported to QIIME2 (Bolyen et al., 2019). Subsequently, forward and reversed sequences were trimmed to a length of 245 and 200 nt, respectively, using the DADA2 plugin (Callahan et al., 2016). The DADA2 analysis produced a quality filtered table of all operational taxonomic units (OTUs), a .fasta file of representative sequences, and a table summarising the denoising statistics. Using the described method, the OTUs generated are equal to exact amplicon sequence variants (ASVs). Following this, the QIIME2 feature-classifier plugin with the extract-reads option was used to extract reads from the Silva database with the silva-138-99-seqs.qza file as input and the forward and reverse PCR primers as parameters. This produced a file of reference sequence reads, which was used as input for the feature-classifier plugin with the fit-classifier-naive-bayes option. The plugin trained a Naive Bayes classifier and produced a classifier file (classifier.qza) as output. The feature-classifier plugin was then used with the classify-sklearn alogrithm using the representative sequences file generated by the DADA2 analysis as input, which produced a table with taxonomic assignment for all OTUs. Mitochondria, chloroplasts, and Eukaryota OTUs were filtered out using the QIIME2 taxa plugin with the filter-table method. The OTU and taxonomy tables were later merged in R. After quality control and removal of singletons, chloroplasts and mitochondria, the total dataset of sediment, water and host bacterial samples consisted of 1549657 sequences and 13711 OTUs. The OTU count table and a fasta file containing all sequences are presented in Supplementary Table 1 and 2, respectively.

**Table 1.**
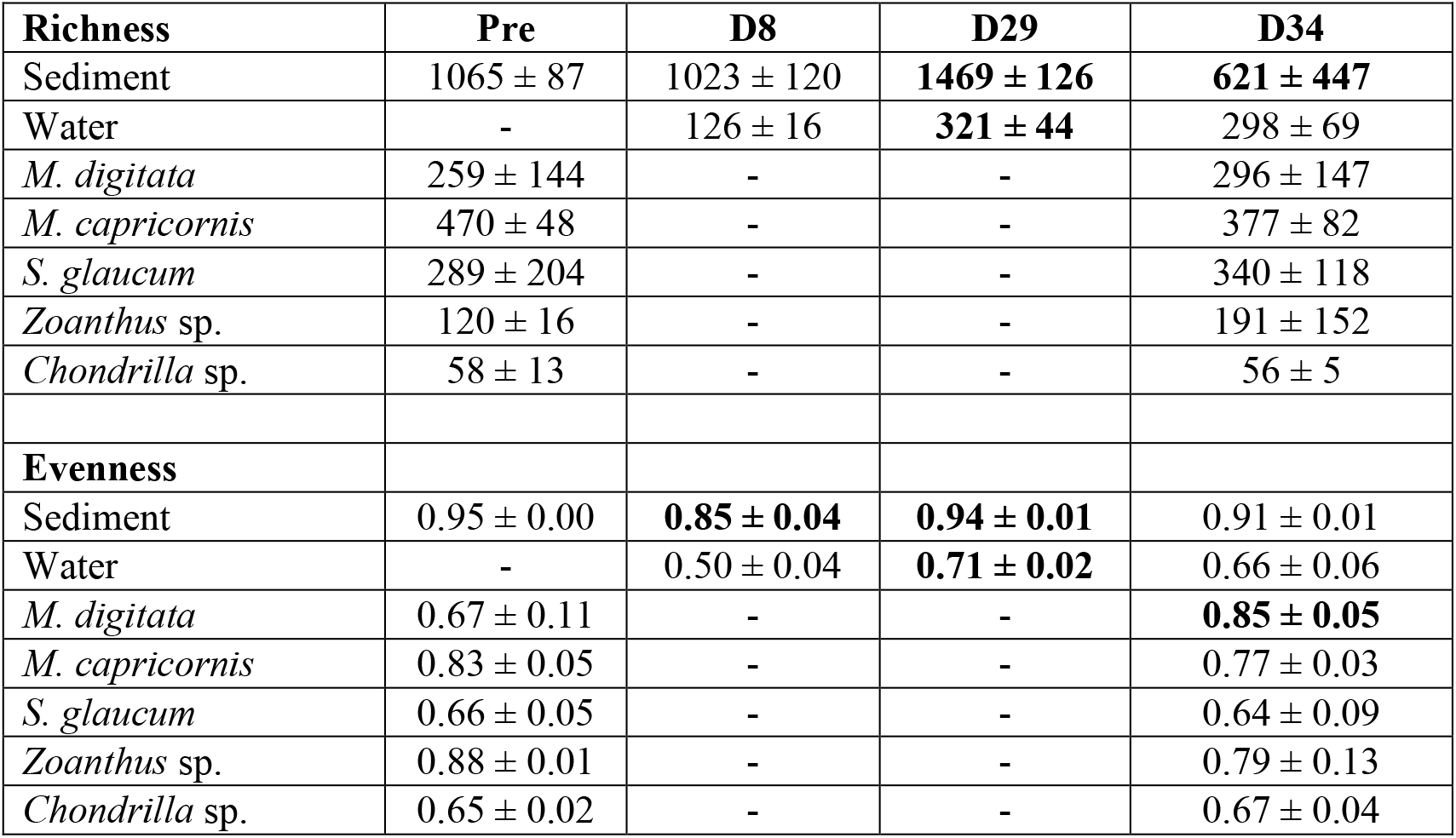
Mean and standard deviations of rarefied richness & evenness. Values which significantly varied from the previous sampling event are indicated in bold (P < 0.004, GLM with emmeans, see supplementary Table 4).

| Richness | Pre | D8 | D29 | D34 |
| --- | --- | --- | --- | --- |
| Sediment | 1065 ± 87 | 1023 ± 120 | **1469 ± 126** | **621 ± 447** |
| Water | - | 126 ± 16 | **321 ± 44** | 298 ± 69 |
| *M. digitata* | 259 ± 144 | - | - | 296 ± 147 |
| *M. capricornis* | 470 ± 48 | - | - | 377 ± 82 |
| *S. glaucum* | 289 ± 204 | - | - | 340 ± 118 |
| *Zoanthus* sp. | 120 ± 16 | - | - | 191 ± 152 |
| *Chondrilla* sp. | 58 ± 13 | - | - | 56 ± 5 |
| Evenness |  |  |  |  |
| Sediment | 0.95 ± 0.00 | **0.85 ± 0.04** | **0.94 ± 0.01** | 0.91 ± 0.01 |
| Water | - | 0.50 ± 0.04 | **0.71 ± 0.02** | 0.66 ± 0.06 |
| *M. digitata* | 0.67 ± 0.11 | - | - | **0.85 ± 0.05** |
| *M. capricornis* | 0.83 ± 0.05 | - | - | 0.77 ± 0.03 |
| *S. glaucum* | 0.66 ± 0.05 | - | - | 0.64 ± 0.09 |
| *Zoanthus* sp. | 0.88 ± 0.01 | - | - | 0.79 ± 0.13 |
| *Chondrilla* sp. | 0.65 ± 0.02 | - | - | 0.67 ± 0.04 |

Subsequently, we removed all OTUs classified as Archaea and OTUs that were unassigned at the phylum level. We also removed OTUs, which occurred in the triple-autoclaved commercial sediment (control for sampling and eDNA contamination) and negative controls used for sequencing (removed OTUs are listed in Supplementary Table 3). OTUs removed following detection in these samples are listed in Supplementary Table 3. Overall, the removed OTUs were assigned to known contaminants, for example, the genera *Ralstonia, Burkholderia-Caballeronia-Paraburkholderia, Reyranella, Bacillus* and *Bradyrhizobium* (Glassing et al., 2016; Salter et al., 2014; Weyrich et al., 2019). After these filtering steps, we removed two samples with low read counts before subsequent analysis; E2G4 (339 sequences) and E2Z1 (1004 sequences). The resulting dataset consisted of 62 samples, 1414326 sequences and 13225 OTUs, also known as ASVs using QIIME2.

#### 2.6.3. Predicted metagenomic analysis

To analyse the putative functional profile of the bacterial communities, we used the Tax4Fun2 library (Wemheuer et al., 2020) in R. Tax4Fun2 predicts the metagenomic content of the samples using the KEGG (Kyoto Encyclopedia of Genes and Genomes) database. Output of Tax4Fun2 consists of a table of functional counts for individual pathways and KEGG orthologs (KOs). Note that because of functional overlap, some KEGG orthologs (KOs) can be represented in multiple pathways. Note that the Tax4Fun2 results as presented are predictive and thus provide information on potential enrichment and putative function as opposed to measuring actual gene presence/expression and function. In a comparative study of PICRUSt (Langille et al., 2013), Tax4Fun (Aßhauer et al., 2015) and Tax4Fun2 using marine seawater and kelp samples, Tax4Fun2 outperformed the other two programs for predictive metagenomic profiling (Wemheuer et al., 2020). Functional profiles, predicted using Tax4Fun2 were, furthermore, highly correlated with functional profiles from the metagenomes of the aforementioned studies. Tax4Fun2 also produces output on the amount of OTUs unused in the prediction for each sample and the amount of sequences unused in the prediction for each sample. This can vary among samples depending on the availability of closely-related, sequenced genomes available in the Ref100NR database.

#### 2.6.4. BLAST

The 50 most abundant OTUs were referenced against the NCBI nucleotide database using NCBI Basic Local Alignment Search Tool (BLAST) (Boratyn et al., 2013). BLAST identifies locally similar regions between sequences, compares sequences to extant databases and assesses the significance of matches; functional and evolutionary relationships can subsequently be inferred. Each BLAST run produces a list of results between the sequence in question (the query sequence), obtained from selected OTUs in the present study, and target sequences in the NCBI nucleotide database.

### 2.7. Statistical analysis

We tested for significant differences among sampling events in SO_4_^2-^and TOC concentrations, and Fv/Fm ratios. Histograms revealed significant deviations from normality. The distributions remained significantly deviant after logarithmic and square root transformations. We, therefore, tested for significant differences in SO_4_^2-^and TOC concentrations, and Fv/Fm ratios among sampling events using a repeated measures permutational analysis of variance with the adonis2() function (Vegan package) in R. In the function, the method was set to “euclidean” and permutations to “999”.

To analyse the bacterial community data, a table containing the OTU counts was imported into R using the read.csv function. The OTU table was used to analyse the impact of the microcosm system on bacterial diversity, composition and higher taxon abundance. Diversity indices were obtained using the rarefy and diversity functions from the vegan package (Oksanen et al., 2019) package in R. Evenness was calculated by dividing Shannon’s H’ by the number of OTUs in each sample. Differences in diversity indices, higher taxon abundances and selected KEGG pathways were investigated using an analysis of deviance (glm function of the R package stats). In the function, we set the family argument to “quasipoisson” (richness) or “quasibinomial” (evenness, higher taxa and KEGG pathways), which is appropriate when analysing proportional data and allowed us to model overdispersion. Using the glm model, we tested for significant variation using the anova() function in R with the F test, which is most appropriate when dispersion is estimated by moments as is the case with quasibinomial fits. We subsequently used the emmeans() function in the emmeans library (Lenth et al., 2020) to perform multiple comparisons of mean abundance among sample events using the false discovery rate (fdr) method in the adjust argument.

Variation in bacterial composition among groups (timepoints and biotopes) was visualised with Principal Coordinates Analysis (PCO). For the PCO, the OTU table was rarefied to the minimum sample size using the rrarefy function of the R package vegan (4818 sequences in the present study). For compositional analyses, the OTU table was transformed using the decostand function in vegan with the method argument set to ‘Hellinger’. With this transformation, the OTU table is adjusted such that subsequent analyses preserve the chosen distance among objects (samples in this case). The OTU table was transformed because of the inherent problems with the Euclidean-based distance metric, which is frequently used in cluster analyses (Legendre and Gallagher, 2001). The Hellinger (Rao, 1995) distance was chosen because it gave very good results in comparison to various distance metrics. In particular, it gave low weights to rare species, was monotonically related to the geographic distance along a model gradient, and reached an asymptote for sites with no species in common. It also produced little ‘horseshoe effect’ or inward folding of sites at opposite ends of the gradient, in ordinations. A distance matrix was subsequently created with the vegdist function in vegan using the Hellinger-transformed OTU table as input and the method argument set to “euclidean”. Subsequently, we used the cmdscale function of the R package stats with the Hellinger transformed distance matrix as input. A permutational anova was performed using the adonis2 function of the R package vegan to test for significant differences in composition among sampling events per biotope (999 permutations). Detailed descriptions of the functions used here can be found in R (e.g., ?cmdscale) and online in reference manuals (http://cran.rproject.org/web/packages/vegan/index.html).

## 3. Results

### 3.1. Physical and (bio)chemical analysis

Water temperatures varied from a minimum of 26.4 ± 0.65°C in the morning to a maximum of 28.3 ± 0.83°C in the afternoon. Dissolved oxygen levels varied over the day from a minimum of 7.83 ± 0.50 mg L^−1^ in the morning to a maximum of 8.60 ± 0.45 mg L^−1^ in the afternoon. In line with variation in oxygen concentrations, pH varied over the day from 8.06 ± 0.04 to 8.17 ± 0.08 (Supplementary Fig. 2). During the course of the experiment, NO3-, NO2-, NH4+ and PO43-concentrations in the water column and pore water remained below detection limit (<0.2, <0.01, <0.15 and <0.05 mg L^−1^, respectively). There was a significant reduction in the concentration of pore water SO42-from 2940.0 ± 16.33 mg L^−1^ at the first measurement to 2520.0 ± 40.82 mg L^−1^ at the end of the experiment (repeated measures PERMANOVA, F2,9 = 18.89, R2 = 0.81, P = 0.035, Supplementary Fig. 2). Although not significant, TOC increased from 1.77 ± 0.18 to 4.97 ± 2.59 mg C L^−1^ at the end of the experiment (repeated measures PERMANOVA, F2,9 = 3.28, R2 = 0.42, P = 0.15, Supplementary Fig. 2).

### 3.2. Coral photosynthetic efficiency

The analysis of the coral photosynthetic efficiency showed that the maximum photosynthetic yield significantly increased from 0.46 ± 0.01 to 0.53 ± 0.01 in *M. digitata* (PERMANOVA: F1, 6 = 148.56; P = 0.031; R2 = 0.96). Fv/Fm ratios increased non-significantly from 0.52 ± 0.01 to 0.54 ± 0.02 in *M. capricornis* (PERMANOVA: F1, 6 = 3.50; P = 0.157; R2 = 0.37) and non-significantly from 0.52 ± 0.04 to 0.59 ± 0.03 in *S. glaucum* (PERMANOVA: F1, 6 = 10.41; P = 0.062; R2 = 0.63); in *Zoanthus* sp. Fv/Fm ratios decreased non-significantly from 0.51 ± 0.02 to 0.44 ± 0.07 (PERMANOVA: F1, 6 = 3.46; P = 0.174; R2 = 0.37).

### 3.3. Bacterial community analysis

#### 3.3.1. Diversity and higher taxonomic composition

The results for OTU richness and evenness for each biotope are summarized in Table 1 and the significance analysis is presented in Supplementary Table 4. Richness and evenness were highest within sediment bacterial communities, but varied among sampling events. After a non-significant increase from 1065 ± 87 at day 8 to 1469 ± 126 at day 29, sediment richness declined significantly from day 29 to 621 ± 447 at day 34. Evenness, in turn, declined significantly from 0.95 ± 0 in Pre samples to 0.85 ± 0.04 at day 8 and subsequently significantly increased from day 8 to 0.94 ± 0.01 at day 29 and remained high (> 0.91). In water, richness and evenness both significantly increased from 126 ± 16 and 0.5 ± 0.04 at day 8 to 321 ± 44 and 0.71 ± 0.02 at day 29 and remained stable at day 34. Richness and evenness did not vary significantly among sampling events in host biotopes, with exception of a significant increase in evenness from 0.67 ± 0.11 in Pre to 0.85 ± 0.05 in day 34 samples in *M. digitata*.

Ordination analysis (Fig. 3) showed that sampling event was a significant predictor of variation in the bacterial community composition of the sediment, water, and host organisms *M. digitata* and *Chondrilla* sp. (PERMANOVA: P<0.05, Supplementary Table 5). In water and sediment, the main axis of variation separated samples according to biotope whereas the second axis separated water and sediment samples collected at day 8 from Pre, day 29 and day 34 samples.

**Fig. 3.**
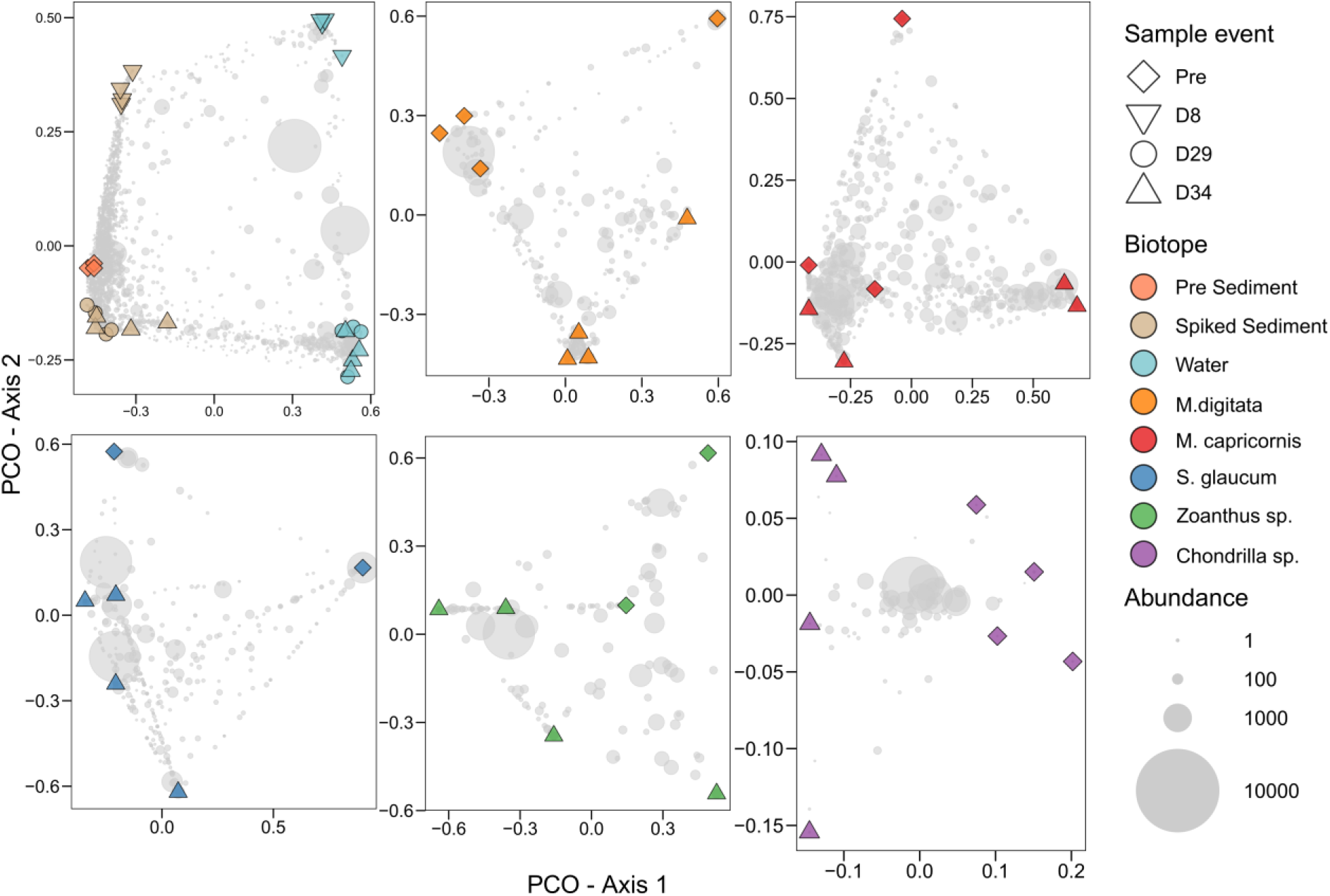
Ordinations showing the first two axes of the principal coordinates analysis (PCO) of prokaryote OTU composition for sediment and water, and each host species separate. Light grey symbols represent operational taxonomic unit (OTU) scores with the symbol size representing their abundance (number of sequence reads). Supplementary Table 5 lists eigenvalues, the total variation explained and results of the PERMANOVA for each ordination.

The most abundant phyla across the complete dataset were Proteobacteria (4156 OTUs, 751894 sequences), Planctomycetota (2214 OTUs, 138421 sequences), Bacteriodota (1634 OTUs, 84631 sequences), Cyanobacteria (221 OTUs, 78311 sequences), Actinobacteriota (485 OTUs, 59621 sequences), Firmicutes (215 OTUs, 54368 sequences), Verrumicrobiota (1112 OTUs, 52766 sequences) and Patescibacteria (490 OTUs, 45003 sequences) (Supplementary Table 6). Of these, Cyanobacteria were particularly abundant in *Chondrilla* sp., whereas Patescibacteria were specifically abundant in water communities (Fig. 4).

**Fig. 4.**
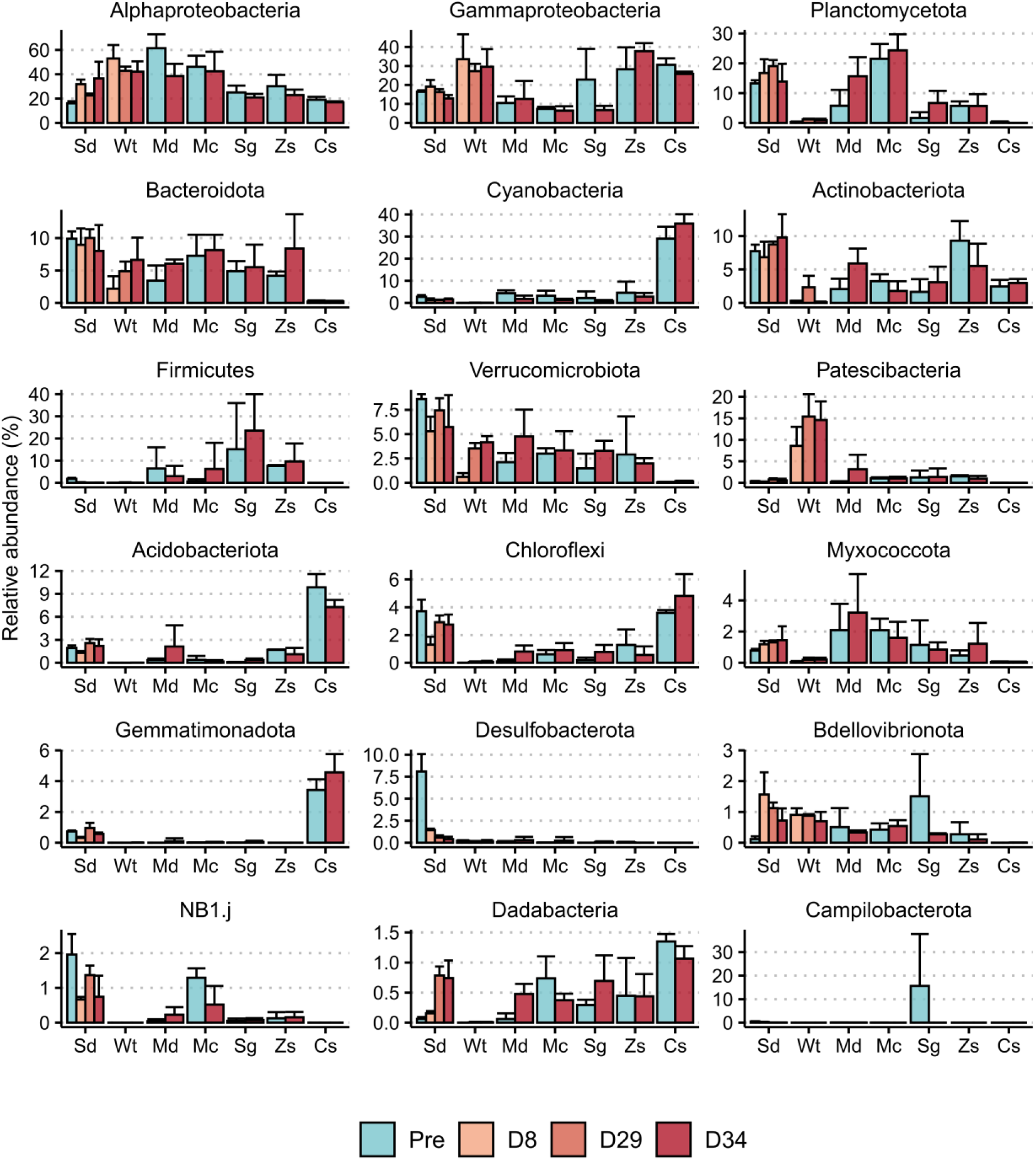
Mean relative abundance of the most abundant proteobacterial classes and phyla of biotopes at the different sample events. Error bars represent standard deviations of the mean. Sd: sediment, Wt: water, Md: M. digitata, Mc: M. capricornis, Sg: S. glaucum, Zs: Zoanthus sp., Cs: Chondrilla sp.

The relative abundances of the most abundant phyla and proteobacterial classes varied significantly among sampling events in water and sediment communities (Fig. 4), whereas no significant differences were observed between sampling events in host organisms (Pre and day 34) (emmeans, Supplementary Table 7). In sediment, Desulfobactereota, Firmicutes and Spirochaetota abundances were significantly (P < 0.0004) lower at day 8, 29 and 34 compared to Pre samples, whereas the abundance of Dadabacteria was significantly higher at day 29 and 34 compared to Pre and day 8 (emmeans, Supplementary Table 7). Chloroflexi abundance first declined significantly from Pre to day 8, increased significantly from day 8 to 29, and did not vary significantly from day 29 to 34, whereas the inverse held for Gammaproteobacteria (emmeans, Supplementary Table 7). In water, the abundance of Verrucomicrobiota and Planctomycetota significantly increased from day 8 to 34 (emmeans, Supplementary Table 7).

#### 3.3.2 Most abundant OTUs and their closest relatives

Of the 50 most abundant OTUs, five were recorded in sediment across all sample events (Fig. 5). These OTUs were assigned to the genera *Ruegeria* (OTUs 5, 25), *Methyloceanibacter* (OTU-36), *Filomicrobium* (OTU-37) and the family Geminicoccaceae (OTU-67). More in depth analysis showed that OTUs 5 and 25 were related to *R. lacuscaerulensis* isolates (100% sequence similarity), whereas the other three OTUs were related to uncultured organisms, previously detected in marine sediment (OTU 67), intertidal outcrops (OTU 37), and the coral *Galaxea fascicularis* (OTU 36) (>99% sequence similarity, Supplementary Table 8). Except for OTU 67, all of these OTUs were also found in water (OTUs 5 and 36) and/or host organisms (OTUs 5, 25, 36, 37) across sampling events.

**Fig. 5.**
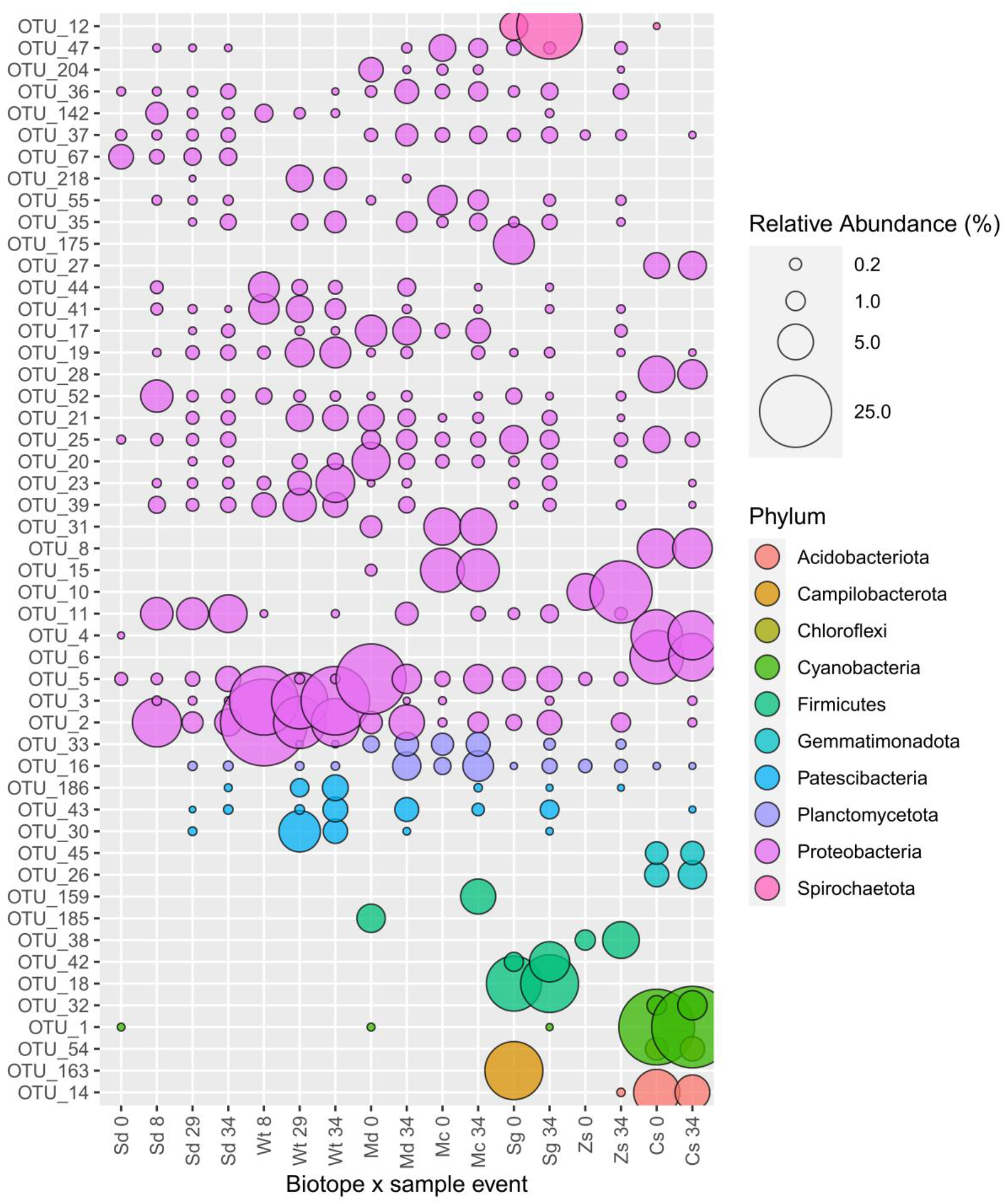
Relative abundance of the 50 most abundant OTUs colour-coded according to bacterial phylum. The circle size of the OTU is proportional to the mean percentage of sequences per biotope and sampling event as indicated by the symbol legend in the bottom right corner of the figure. The y-axis numbers show the OTU id number. Sd: sediment, Wt: water, Md: M. digitata, Mc: M. capricornis, Sg: S. glaucum, Zs: Zoanthus sp., Cs: Chondrilla sp.

Ten OTUs, assigned to the Alpha- and Gammaproteobacteria (OTUs 2, 11, 19, 23, 39, 41, 52, 55, 3 and 142) were consistently abundant in sediment at day 8, 29 and 34 but were not detected in Pre sediment. They also occurred in water and invertebrate hosts at various sampling events. OTUs 2, 11, 52 and 142 were related (100% sequence similarity) to isolates classified as *Tritonibacter litoralis, Leisingera* sp., *Ruegeria* sp. strain and *Alteromonas* sp. detected in seawater, a coral, marine sediments and estuarine soil, respectively. OTUs 3, 19, 23, 39, 41 and 55 were related to (> 97% sequence similarity) uncultured organisms detected in seawater, the coral *Fungia granulosa*, the coral *Alcyonium digitatum*, the sponge *Haliclona* sp., endolithic communities and marine sediment, respectively. Three OTUs (30, 43 and 186), assigned to the classes Gracilibacteria and ABY1, were not detected in sediment and host Pre samples, but were abundant across water and sediment at day 29 and 34 and associated with several invertebrate hosts at day 34. Additionally, eleven of the 50 most abundant OTUs showed a biotope specific association (only recorded in a single biotope). For example, OTU 42 (*Entoplasmatales*) was only recorded in *S. glaucum*, OTU-10 (*Endozoicomonas*) in *Zoanthus sp*., and OTUs 26 (BD2-11 terrestrial group) and 32 (Cyanobiaceae) in *Chondrilla sp*.. These OTUs were similar to uncultured organisms previously detected in the sea anemone *Nematostella vectensis*, a deep-sea octocoral, and the sponges *Xestospongia muta* and *Haliclona* sp., respectively (Supplementary Table 8).

#### 3.3.3. Predicted functional analysis

In sediment, the predicted relative abundances of 7 of the 12 tested KEGG categories remained relatively stable across all sampling events. Variation was observed in the nitrogen metabolism category, which significantly declined from Pre to day 8, but increased from day 8 to 29 and 34 to similar values found in Pre sediments (P < 0.0006, Fig. 6 and Supplementary Table 9). Additionally, predicted gene count abundances of the cAMP signalling and two-component system categories significantly (P < 0.0006) decreased, and the carbon metabolism and quorum sensing categories significantly increased from Pre to day 8 and thereafter remained relatively stable from day 8 to 34 (emmeans, Supplementary Table 9). In water, 6 of the categories remained stable. Variation was observed among the carbon metabolism, secondary metabolites and antibiotic biosynthesis categories, which increased from day 8 to 34, whereas the quorum sensing category decreased (emmeans, Supplementary Table 9). The terpenoid backbone biosynthesis category increased from day 8 to 29, but there was no significant difference between day 8 and 34. The cAMP signalling category significantly increased from day 8 to 29 and subsequently significantly decreased from day 29 to 34 (emmeans, Supplementary Table 9). In host biotopes, none of the functional categories investigated differed significantly between sampling events.

**Fig. 6.**
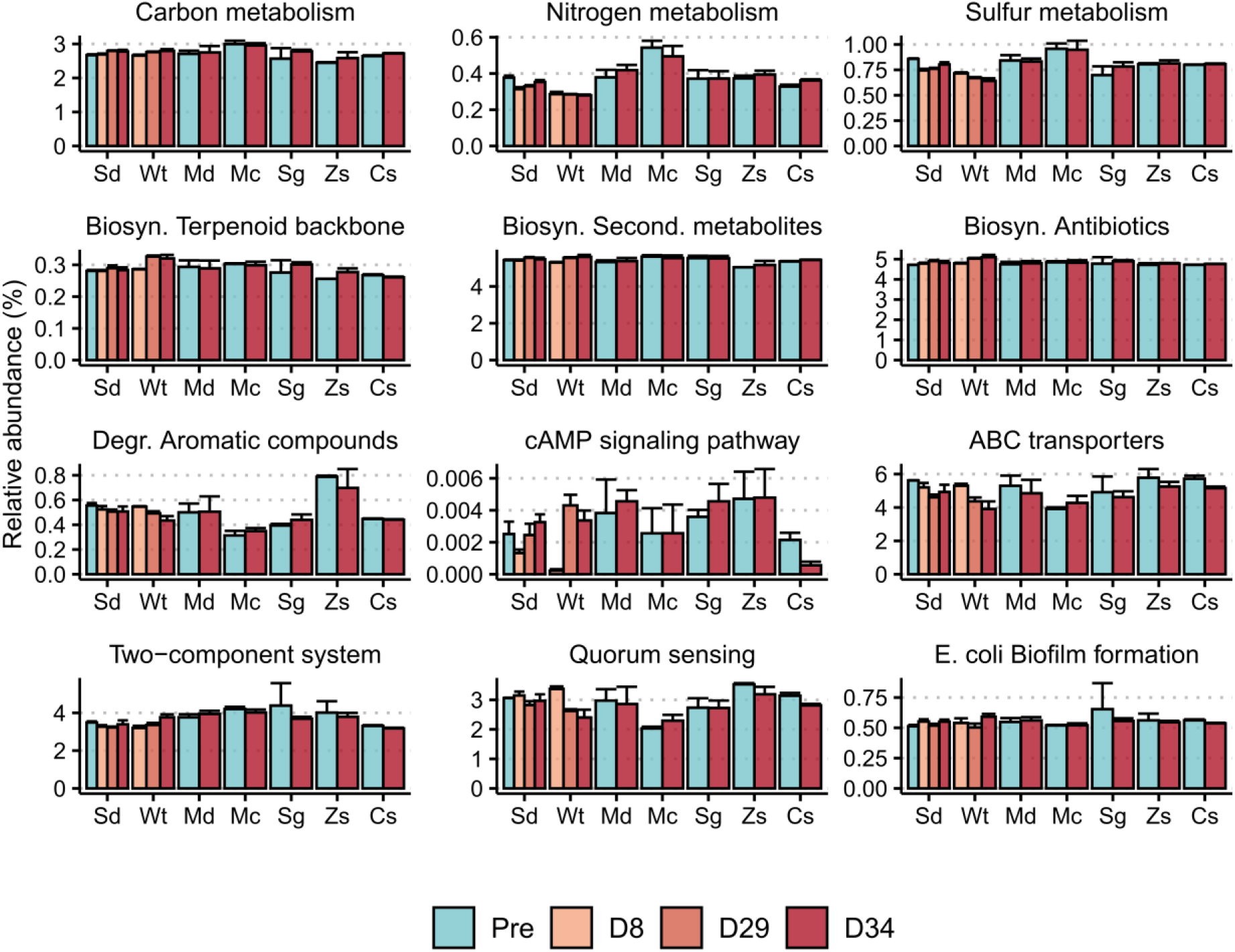
Predicted mean relative gene count abundance of the KEGG level 1 categories of biotopes at the different sample events. Error bars represent standard deviations of the mean. Sd: sediment, Wt: water, Md: M. digitata, Mc: M. capricornis, Sg: S. glaucum, Zs: Zoanthus sp., Cs: Chondrilla sp.

## 4. Discussion

The current study aimed to develop and validate an ELSS model system for studies on the impact of ultraviolet B radiation and temperature on coral reef microbial communities. For this goal, we assessed bacterial community composition (sediment, water, and host-associated communities), coral photosynthetic activity, and physicochemical parameters (e.g., salinity, pH, temperature, PAR and UV light, inorganic nutrients and DOC) within the ELSS and compared the results with natural coral reef environments.

### 4.1. Physiochemical parameters resembled natural conditions

Average values of the water temperature, dissolved oxygen, pH and salinity in the microcosms were similar to conditions at shallow coral reef sites (Bainbridge, 2017; DeCarlo et al., 2017; Guadayol et al., 2014). Daily fluctuations were observed among water temperatures (increased by 2 degrees during the course of the day), oxygen and pH (higher in the morning and lower in the afternoon). At reefs, similar temperature fluctuations have been observed due to sun exposure (Bainbridge, 2017; DeCarlo et al., 2017), whereas oxygen and pH can fluctuate as a result of net-photosynthesis during the day and respiration at night (Guadayol et al., 2014).

Inorganic nutrients in the water column and sediment pore water (NO3-, NO2-, NH4+ and PO43-) remained low (<0.2, <0.01, <0.15 and <0.05 mg L^−1^, respectively) throughout the experiment. Inorganic nutrient concentrations below these values were previously observed in pacific coral reefs as well (Silbiger et al., 2018). Porewater sulphate concentrations of surface sediments have been observed to vary between 2300 - 2880 mg L^−1^ (Alongi, 1995; Werner et al., 2006). At the first timepoint, sulphate was relatively high (2940.0 ± 16.33 mg L^−1^) in the microcosms, but fell to levels previously observed by Alongi (1995) and Werner et al. (2006) in coral reef environments. Although we detected relatively low DOC in the ELSS at the beginning of the experiment (1.77 ± 0.18 mg C L^−1^), DOC concentrations increased to 4.97 ± 2.59 mg C L^−1^ at day 34. These values were in the range of DOC concentrations (∼ 3 to 8 mg C L^−1^) in surface sediment pore water at Great Barrier Reef sites (measured using 0.4 μm membrane filters, Lourey et al. 2001). It should also be noted that the membrane pore size of our samplers was 0.6 μm, which included a slightly larger fraction of total organic matter compared to the measurements taken at the Great Barrier Reef.

### 4.2. Photosynthetic efficiency of coral endosymbionts

The photosynthetic efficiency of *M. capricornis, S. glaucum* and *Zoanthus* sp. did not change significantly following 34 days in the microcosms, which indicates that the photo-physiology of these organisms was stable and showed no signs of stress. The photosynthetic efficiency of *M. digitata*, however, was significantly increased at day 34 compared to Pre conditions. The increased photosynthetic efficiency can indicate a certain level of photo-acclimation of the photosynthetic endosymbionts of *M. digitata* to the light conditions in the ELSS (Roth, 2014; Roth et al., 2010).

### 4.3. ELSS bacterial communities

#### 4.3.1. Community composition and diversity

In line with previous studies, our analysis showed that bacterial evenness and richness were lowest in the sponge *Chondrilla* sp. and highest in the sediment (Cleary et al. 2019, 2020). In both water and sediment, there were significant shifts in bacterial evenness, and composition from day 8 to 29, but not from day 29 to 34, suggesting microbiome stabilisation. Additionally, sediment bacterial evenness declined significantly from Pre to day 8 and subsequently increased from day 8 to 29 and 34. The same applied for the bacterial community structure (PCO) with day 8 deviating significantly from both the Pre and day 29 and 34 sampling events. Overall, our results indicate a return to natural-type conditions by day 29. Sediment bacterial richness, however, significantly declined from day 29 to 34. Previously, Coelho et al. (2013) also observed lowered bacterial richness in microcosms compared to natural sediment. They attributed this to the more stable and less heterogeneous conditions in microcosms. Bacterial richness and evenness in the present study fell within ranges observed at natural reef sites for all host-associated biotopes with the exception of *Zoanthus* sp., where relatively low richness values were observed compared to previous findings (Cai et al., 2018; O’Brien et al., 2020; Shore-Maggio et al., 2015; Sun et al., 2014). Richness and evenness at specific time points varied more among individuals of hard and soft coral biotopes than among individuals of the sponge *Chondrilla* sp., in line with previous in situ comparisons of corals and sponges (O’Brien et al., 2020). Evenness only differed significantly between sampling events (Pre and day 34) in *M. digitata*, increasing substantially from Pre to day 34. Bacterial composition, in turn, differed significantly between Pre and day 34 for *M. digitata* and *Chondrilla* sp., but not for the other host-associated biotopes. This finding suggests that the bacterial communities of *M. digitata* and *Chondrilla* sp. are able to adapt to changing environmental conditions as exemplified in the present case by transplantation between aquaria. Previous studies have shown that the bacterial communities of certain coral species change along with shifting environmental conditions, a trait, which may be part of the process of host adaptation to new environmental conditions (Lurgi et al., 2019; Vanwonterghem and Webster, 2020; Ziegler et al., 2019).

#### 4.3.2. Bacterial higher taxonomic composition

Dominant classes and phyla observed in the present study were similar to those observed *in situ* in several coral reef studies (Cleary et al., 2019b, 2019a; O’Brien et al., 2020; Sun et al., 2014). Across biotopes, the most abundant higher taxa consisted of Alpha- and Gammaproteobacteria, Bacteriodota, Verrucomicrobiota (with exception of sponges) and Planctomycetota (with the exception of water and sponges). Firmicutes were most abundant in corals, whereas Cyanobacteria, Acidobacteriota and Gemmatimonadota were most abundant in sponges and Patescibacteria in water. In sediment and water, the abundance of several higher taxonomic groups differed significantly among sampling events, whereas no significant variation was observed among host organisms. In sediment, the relative abundance of the phylum Desulfobactereota was significantly higher in Pre compared to day 8, 29 and 34. Bacterial members of this phylum were mainly assigned to the orders Desulfobulbales, Desulfobacterales and Sva1033. These orders include a variety of presumably, anearobic sulfate-reducing bacteria, which are widespread and abundant in marine sediments where they oxidize organic matter by reducing sulfate to sulfide under anoxic conditions (Begmatov et al., 2021; Rabus et al., 2015; Zoppini et al., 2020). This finding indicates that the microcosm sediment layers may have been exposed to higher oxygen concentrations than sediment retrieved from coral reefs, thereby favouring aerobic (e.g., Rhodobacteraceae members) groups. The abundance of the phylum Chloroflexi was significantly lower in day 8 compared to Pre, but increased to abundances similar to Pre at later timepoints. Although not significant, the same pattern was observed for Acidobacteriota and Gemmatimonadota, and indicates recolonization by these bacterial groups after the initial disturbance and a return to natural-type conditions. In water, the relative abundance of Verrucomicrobiota and Planctomycetota increased from day 8 to day 29 and day 34. Members of these bacterial phyla are often observed within marine bacterioplankton communities where they have been associated with microalgae blooms (Orellana et al., 2022; Wiegand et al., 2018). Recently, genomic and proteomic data indicated that small, coccoid, free-living Verrucomicrobiota specialise in the degradation of fucoidan-like substrates during spring algal blooms in the North Sea (Orellana et al., 2022).

#### 4.3.3. Most abundant OTUs and closely related organisms

In sediment, abundant OTUs were assigned to the genera *Ruegeria* (family Rhodeobacteraceae), *Methyloceanibacter* (family Methyloligellaceae) and *Filomicrobium* (family Hyphomicrobiaceae). Members of these families were previously found to be abundant in marine sediment and are known to play an important role in nutrient remineralization (Begmatov et al., 2021; Dong et al., 2022; Lin et al., 2022; Vekeman et al., 2016). Recently, *Ruegeria* strains isolated from marine sediment were observed to utilize various organic and inorganic compounds, oxidize sulfur, and perform complete denitrification by reducing nitrate to molecular nitrogen (Lin et al., 2022). The Methyloligellaceae and Hyphomicrobiaceae families include bacterial species, which are able to reduce one-carbon compounds (methylotrophs) and oxidize methane under aerobic conditions (Vekeman et al., 2016). Additionally, *Methyloceanibacter* members may be involved in nitrogen cycling (Begmatov et al., 2021; Vekeman et al., 2016).

OTU-2, assigned to the Rhodobacteraceae, was particularly abundant in water and had 100% sequence similarity to an organism identified as *Tritonibacter litoralis* isolated from coastal surface seawater (Li et al., 2021). *Tritonibacter* members are globally distributed and primarily found in surface waters (Brinkhoff et al., 2008). They are also often found in association with seaweeds due to their ability to metabolize dimethylsulfoniopropionate (DMSP) (Brinkhoff et al., 2008; Li et al., 2021). Moreover, certain *Tritonibacter* strains (producers of of the antibacterial compound Tropodithietic acid) can inhibit the growth of fast-growing bacteria and fish pathogens (Henriksen et al., 2022). OTU 33, assigned to the Pirellulaceae, was abundant in all cnidarian biotopes in the present study, but absent from other biotopes. Members of this family have previously been detected in deep-sea octocorals and sponges and are believed to play a role in nitrogen cycling (Kellogg et al., 2016; Mohamed et al., 2010). OTU-10, assigned to the genus *Endozoicomonas*, was only recorded in *Zoanthus* sp.. *Endozoicomonas* spp. have been recorded across a range of hosts including corals. Previous studies have suggested that they play important roles in host nutrient cycling and influence microbiome structure (Neave et al., 2016; Pogoreutz et al., 2022). OTUs 1 and 32, assigned to the Ca. *Synechococcus spongiarum*, were specifically enriched in *Chondrilla* sp.. Members of this genus have been previously detected in 28 sponge species and may play a role in carbon assimilation and transfer to their sponge host (Burgsdorf et al., 2022, 2015). OTUs 18 and 42, assigned to the Ca. *Hepatoplasma* (Tenericutes) were only detected in *S. glaucum*. Members of this genus have been suggested to play roles in the degradation of complex organic molecules in the digestive tracts of terrestrial and marine isopods (Bouchon et al., 2016). Notably, three OTUs assigned to the phylum Patescibacteria (OTUs 30, 43, 186) were found across a variety of biotopes from day 29 onward, but were not detected in natural sediment or host-associated bacterial communities sampled prior to addition to the ELSS (Pre). Patescibacteria currently lack isolated representatives but have been detected in marine surface water (Rahlff et al., 2020), sediment (Demko et al., 2021) and reef sponges (Robbins et al., 2021). Members of this phylum are characterized by reduced cell and genome sizes with limited metabolic capacities (Brown et al., 2015). It has been suggested that they live in symbiosis with other bacteria (Lemos et al., 2019) and presumably play a role in oceanic carbon cycling, a finding substantiated by the detection of genes required for carbon fixation in nano-sized patescibacterial members (Lannes et al., 2019).

#### 4.3.4. Putative functional profiles of bacterial communities

Despite of some variation among biotopes, the predicted functional profiles (e.g., degradation of aromatic compounds, nutrient metabolism, and biosynthesis of secondary metabolites and antibiotics) of bacterial communities in water, sediment and host biotopes were relatively stable at different sampling events. These results suggest that the environmental conditions of the ELSS developed in this study did not have significant effects on the community homeostasis. The predicted gene counts of the cAMP signalling and two-component system categories were, however, significantly lower at all time points from day 8 onward in the sediment biotope compared to Pre, whereas the reverse held for the quorum sensing category. The two-component system and cAMP signalling categories help bacteria to adapt to changing environmental conditions (Ohmori et al., 2009; You et al., 2013; Zschiedrich et al., 2016). Quorum sensing, in turn, allows groups of bacteria to synchronously alter gene expression in response to changes in population density and environmental cues (Mukherjee and Bassler, 2019). Quorum sensing has been predicted to give bacteria a competitive advantage under nutrient limiting conditions (Schluter et al., 2016). Previous studies have shown that two-component system category was enriched in members of the phylum Desulfobacterota (Alm et al., 2006), whereas, the quorum sensing category was enriched in members of the class Alphaproteobacteria (Armes and Buchan, 2021; Cude and Buchan, 2013; Zan et al., 2014). These findings suggest an association between higher taxonomic abundance and predicted function.

## 5. Conclusion

The current study described and validated an ELSS model system to assess the effects of UVB and temperature on host and free-living coral reef bacterial communities. The physical and chemical conditions and bacterial communities of the ELSS were similar to those of coral reef ecosystems. Sediment bacterial diversity and composition were more similar between Pre and day 29 and day 34 than between Pre and day 8 suggesting a return to natural-type community diversity. By changing the pre-sets of our ELSS values with respect to temperature and UV, it is possible to simulate several scenarios of global climate change, such as the predicted increase in temperature and UVB.

Temperature can be regulated using water-baths, which allow temperature control in blocks of four microcosms. The transparent polyester films that blocked UVB radiation in our validation trial can be removed from individual microcosms, thus enabling full randomisation. Additionally, irradiance values can be manipulated by varying lamp intensity and time of operation. The modular design of the ELSS developed here offers a multitude of statistically robust experimental designs. In this way, the system enables researchers to establish cause–effect relationships of the individual and interactive effects of a suite of environmental conditions on marine bacterial communities.

## CRediT authorship contribution statement

**T.M Stuij**: Methodology, Investigation, Data analysis, Writing –original draft, review & editing; **D.F.R. Cleary**: Supervision, Conceptualization, Data analysis, Writing – original draft, review & editing; **R.J.M. Rocha**: Resources, Conceptualization, Writing – review & editing; **A.R.M. Polonia**: Investigation, Writing – review & editing; **D.A.M. Silva**: Conceptualization, Investigation, Writing – review & editing; **J.C. Frommlet**: Resources, Writing – review & editing; **A. Louvado**: Conceptualization, Methodology, Investigation, Writing – review & editing; **Y. M. Huang**: Resources, Writing – review & editing; **N. Van der Windt**: Investigation, Writing – review & editing; **N.J. De Voogd**: Supervision, Conceptualization, Resources, Writing – review & editing; **N.C.M. Gomes**: Supervision, Conceptualization, Resources, Methodology, Writing – original draft, review & editing.

## Data availability

Sequences generated in this study can be downloaded from the NCBI Sequence Read Archive under the BioProject accession number PRJNA904682.

## Acknowledgements

We are grateful to F. Coelho and V. Oliveira for their help during lab work and FOLEX COATING GMBH in Germany for providing samples of their specialized polyester film. The research was conducted at the Laboratory of Molecular Studies and Marine Environment, situated at the Centre for Environmental and Marine Studies (CESAM), University of Aveiro, Portugal.

