## Supplementary material for "Development and validation of an experimental life support system to study the impact of ultraviolet B radiation and temperature on coral reef microbial communities"

Postal address:
Newton Gomes
Centre for Environmental and Marine Studies (CESAM)
Second floor, building 2
Campus Universitário Santiago,
3810-193 Aveiro, Portugal

### Coral Fragmentation

Of the five model species, larger individuals were fragmented into 2-3 cm pieces and glued (acrylite) or stitched (*Sacrophyton glaucum*) to small carbonate sand stones (3 cm in diameter, 1 cm thickness). To recover from the fragmentation process, coral fragments were kept in coral reef aquaria (Rocha et al. 2015) for four months before transplantation to the ELSS.

Rocha, R.J.M., Bontas, B., Cartaxana, P., Leal, M.C., Ferreira, J.M., Rosa, R., Serôdio, J., Calado, R., 2015. Development of a Standardized Modular System for Experimental Coral Culture. J. World Aquac. Soc. 46, 235–251. https://doi.org/10.1111/jwas.12186

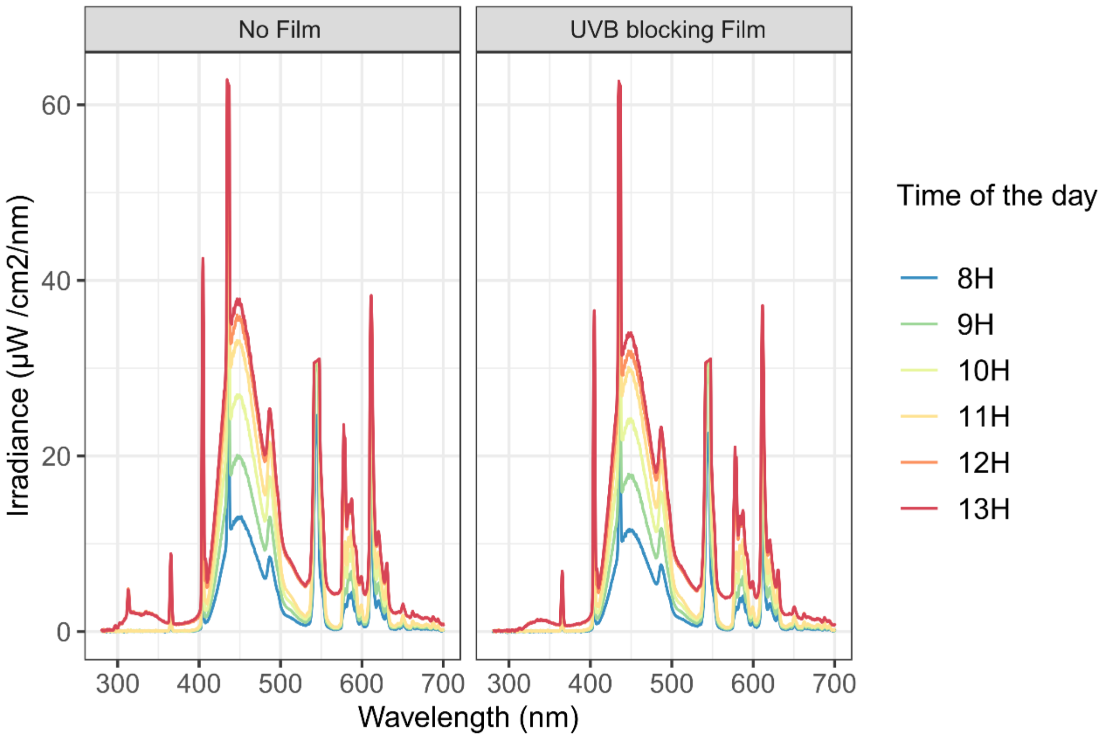


Supplementary Fig. 1 Irradiance spectrum of the fluorescent lamps measured at the point where the light would hit the water surface with and without the transparent polyester film (Folanorm SF-AS, Folex coating, Köln, Germany). Light intensity increased from 8.00 in the morning to 13.00 in the afternoon.

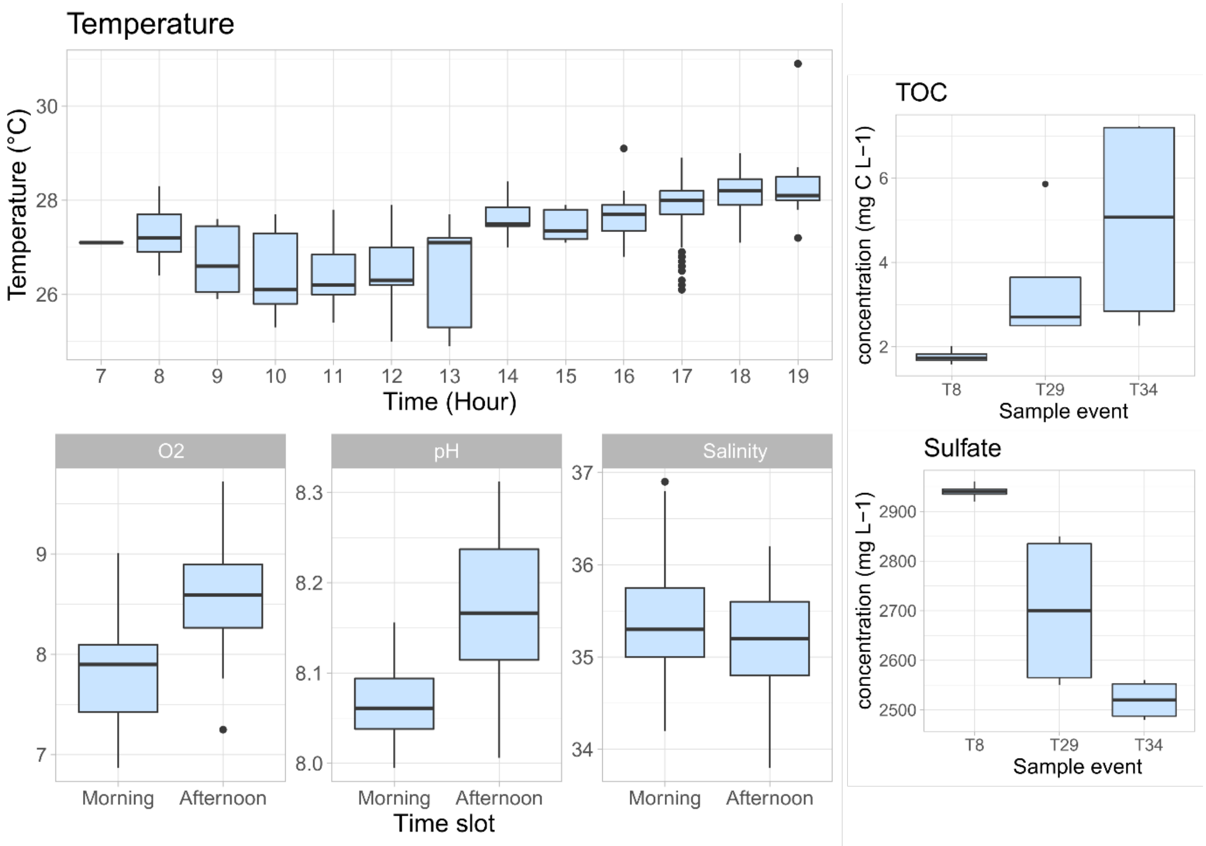


Supplementary Fig. 2 Physical and chemical parameters measured at different timepoints during the experiment. Temperature (hourly), oxygen, pH and salinity (two times per day) were measured in the water column. TOC and Sulphate were measured in the sediment porewater at day 8, 29, and 34.
